# Evolution of behavioral resistance in host-pathogen systems

**DOI:** 10.1101/2020.05.17.100453

**Authors:** Caroline R. Amoroso, Janis Antonovics

**Author notes:** Correspondence CRA, JA.

## Abstract

Behavioral resistance to parasites is widespread in animals, yet little is known about the evolutionary dynamics that have shaped these strategies. We show that theory developed for the evolution of physiological parasite resistance can only be applied to behavioral resistance under limited circumstances. We find that accounting explicitly for the behavioral processes, including the detectability of infected individuals, leads to novel dynamics that are strongly dependent on the nature of the costs and benefits of social interactions. As with physiological resistance, the evolutionary dynamics can also lead to mixed strategies that balance the costs of disease risk and the benefits of social interaction, with implications for understanding avoidance strategies in human disease outbreaks.

## Introduction

Hosts resist parasites using diverse mechanisms, with broad implications for host-parasite coevolution [1–4]. Previous theoretical models of resistance evolution have largely focused on physiological or biochemical resistance [1,5–7]. Yet resistance against parasites can also take the form of behavioral traits [3,8,9] such as direct avoidance or “disgust” in response to diseased individuals [10,11] or general avoidance of interactions with other individuals, which in a human context is now termed “social distancing.” Ecologists have long recognized that social behaviors can both facilitate and prevent transmission [12–14]. Here, we ask whether the ecological and evolutionary dynamics of behavioral defenses against parasites operate according to similar principles as physiological defenses.

Host behavior is implicit in classical models of microparasite transmission as a component of the parameter β, the transmission coefficient[15,16]. β is a composite of multiple factors [17], including the contact rate between hosts, which is a function of host behavior, and the per contact transmission probability, which is a function of the infected host’s infectiousness and the susceptible host’s physiology [18]. Models of resistance evolution typically vary the physiological resistance of the susceptible host [5,19]. Nevertheless, variation in avoidance of infected conspecifics exists across and within species: e.g., crustaceans [20], birds [21,22], and primates [23], including humans [10]. Despite the broad diversity of these behaviors [8,24,25], behavioral resistance has rarely been examined explicitly in theoretical contexts [26,27]. It remains unknown whether evolutionary dynamics of behavioral resistance follow the same patterns as physiological resistance. Previous theoretical research on physiological resistance has shown that susceptible and resistant individuals can coexist in the presence of a disease when resistance carries a direct physiological cost [5,7,28], but in social species, lost interactions with others as function of avoiding disease could constitute a social cost. Models have not yet considered how such costs might influence resistance evolution.

Here, we develop a theoretical model of a disease transmitted in a social context, through direct contact or aerosol, and investigate the evolution of behavioral resistance under several assumptions about behavioral processes and cost-benefit trade-offs. We show with this heuristic model that behavioral resistance can result in evolutionary dynamics that differ from physiological resistance, depending on the specificity of behavioral responses to diseased conspecifics and the nature of the costs and benefits of sociality.

## The Model

We model social behavior in a population of individuals that enter into groups or remain singletons. Let S be the number of singletons and G the number of groups of size T. Thus, the total population size in a given time-step is the sum of singletons and individuals in groups, *N* = *S* + *TG*. (We provide full derivations of subsequent equations in the Supplementary Information, S1). We assume group formation occurs rapidly, reaching equilibrium within each time-step, prior to transmission, birth, and death. Once groups are formed, disease transmission is only possible within groups. We assume a large population and deterministic dynamics.

### Model Structure

#### Group Formation

The frequency of groups depends on the group encounter rate, ρ, and group dissociation rate, ν. We model the simplest case: pair formation (*T* = 2). Pair formation has been studied in the context of mating and marriage, and represents a complex problem of sampling without replacement [29,30]. Following previous work [31], we considered two forms of encounter. First, singletons could encounter one another at a constant frequency, independent of their density, as would occur when individuals seek others out to form associations. Second, singletons could encounter others randomly, such that encounters occur at a higher rate at greater densities. These two types of group formation have parallels with frequency-dependent and density-dependent disease transmission processes [31]. Given that the two types of encounter gave qualitatively similar results, in the main text we present only the frequency-dependent case (density-dependent results are in Supplementary Information, S2). The differential equations for number of groups and singletons are 

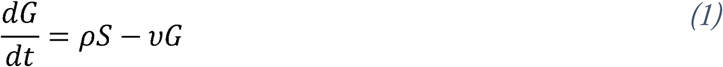

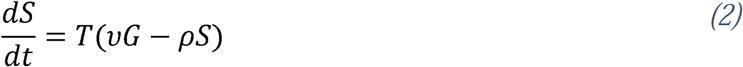

Within a time-step, when pairs form, the total population size (*N*) is fixed. At equilibrium, the ratio of groups to singletons, 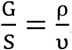. Converting to a frequency, the equilibrium number of groups is 

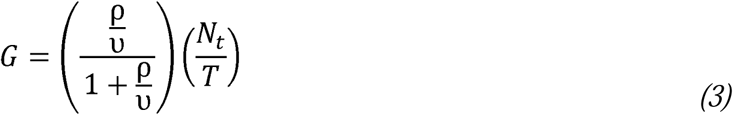

#### Behavioral Resistance

We compare two types of behavioral resistance: specific avoidance of diseased individuals and general avoidance of all associations. For specific avoidance, a healthy individual can detect and avoid pairing only with infected individuals by a factor *ϕ*. For general avoidance, a healthy individual encounters all others at a reduced rate (*ρ* − *a*).

#### Resistance Costs

Physiological resistance is usually assumed to carry some cost that results in reduced fitness in the absence of the parasite [5,28]. We assume behavioral resistance can have two types of cost. Costs of avoidance may be fixed, in that they are incurred regardless of whether avoidance is carried out; for example, a less active genotype could have fewer social encounters, but also reduced feeding. The cost reduces births by *c* relative to the birthrate of non-avoiding individuals, *b*,. Alternatively, sociality could be beneficial, such that costs of avoidance may only be instantiated when the individual avoids being in a group. We examine the case in which reproduction increases as a direct, linear function of the frequency at which each type pairs (see Supplementary Information S1: SE19-SE20).

### Model Implementation

#### Dynamics with No Evolution

We first examine how the equilibrium frequency of individuals in pairs and disease dynamics vary across a range of general and specific avoidance parameters (*ϕ* and *a*) when all individuals avoid disease. We derive how *R*_*0*_ depends on the equilibrium frequency of pairs.

#### Evolution of Behavioral Resistance

To understand the evolution of behavioral resistance we use the one-locus, two allele dynamical framework developed for physiological resistance evolution [5]. In this system, *X*_l_ and *X*_2_ represent two haploid genotypes that differ in their resistance, with *X*_2_ avoiding disease. *X*_l_ and *X*_2_ are equivalent in their transmission once infected and are pooled into one class of diseased individuals, *Y*. We assume that once an individual is diseased, it no longer avoids others. If we assume instead that individuals retain their avoidance once infected, it can be shown that the results are identical for frequency-dependent pair formation, whereas for general avoidance under density-dependent pair formation, Y1 and Y2 need to be distinguished; however, the results are qualitatively equivalent (Supplementary Information S1 and S2).

Transmission occurs at rate *δ* from infected (*Y*) individuals to *X*_l_ or *X*_2_ when they are in a pair. We assume the disease is sterilizing, i.e. diseased individuals do not reproduce. We impose density dependence on birth rate of the healthy individuals because without a numerical (i.e.ecological) feedback, the system does not reach stable equilibrium [28]. We represent background mortality as *μ*. These processes are represented by 

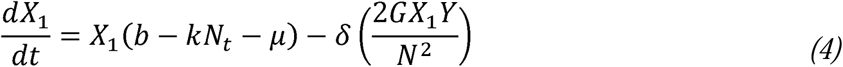

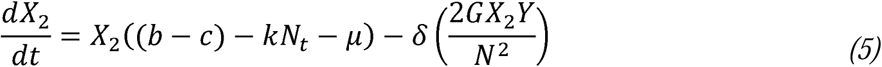

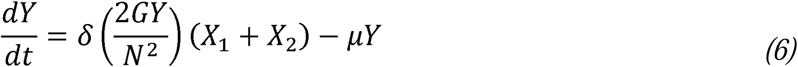

The process of pair formation is nested within each time-step, such that N does not change during pair formation (*N* = + 2*G*), but changes each time-step due to changes in numbers of *X*_l_, *X*_2_, and *Y* individuals.

We obtained equilibria using the differential equation solver (function “ode” Runge-Kutta “rk4” method) from the R package *deSolve* [32,33], and confirmed the stability of the equilibria by perturbation of initial values above and below equilibria.

## Results

### Dynamics with No Evolution

It can be shown that if all individuals are in pairs, i.e. in contact, then the dynamics of disease with pair formation are identical to the physiological resistance model (Supplementary Information S1). We first examined the effect of the different avoidance strategies on the equilibrium frequency of individuals in pairs, prevalence of disease, and *R*_0_ when only *X*_2_, the avoiding genotype, was present (Fig. 1).

**Figure 1.**
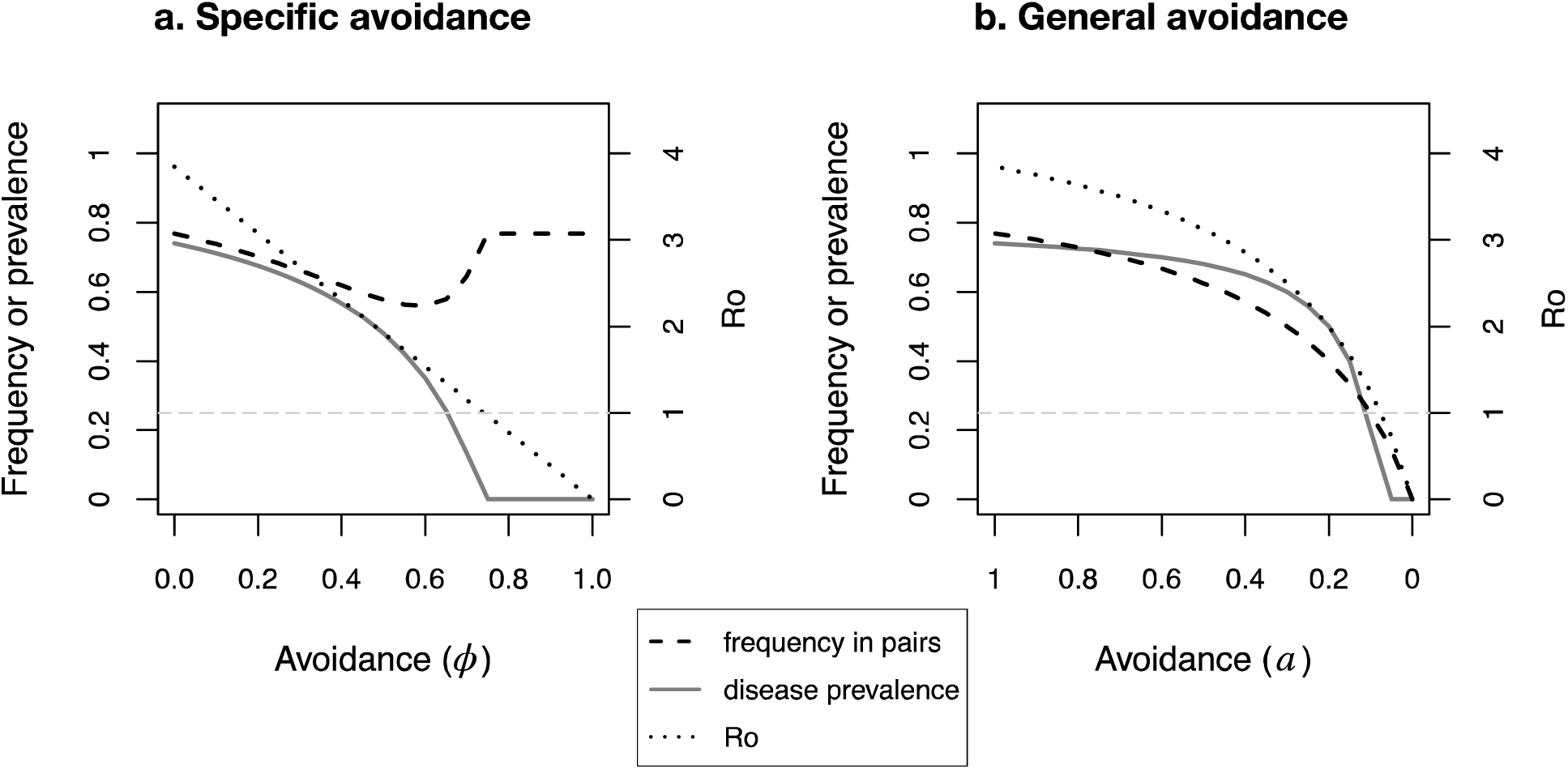
Pairing and disease dynamics at equilibrium when only the avoiding genotype *X*_2_ is present in the population under different avoidance strategies. The light gray horizontal dotted line represents the basic reproductive number *R*_0_ = 1, below which the disease cannot persist in the population, and above which sustained transmission is possible. *b* = 1, *μ* = 0.2, *δ* = 1, *ρ* = 1, *v* = 0.3, *k* = 0.01.

The basic reproductive number of the parasite, 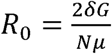, is equivalent to canonical formulations for *R*_0_ for frequency dependent transmission, taking into account the frequency of groups within which transmission occurs. Increased specific avoidance of infected individuals is highly effective at reducing prevalence of the disease, and also results in a decrease in the frequency of individuals in pairs (Fig. 1a). However, at high levels of specific avoidance the frequency of individuals in pairs increases again, because few infected *Y* individuals remain for the *X*_2_ individuals to avoid. With further avoidance, *R*_0_ falls below 1, prevalence drops to 0, and pair formation is only among healthy individuals. Thus, at high levels of specific avoidance, hosts can successfully extirpate the disease from the population while maintaining their social structure.

General avoidance also reduces *R*_0_ and prevalence, but if per-contact transmission rate (δ) is high, avoidance of pairing must be nearly complete to reduce *R*_0_ below the threshold of 1 (Fig. 1b). Therefore, if hosts cannot detect infection in conspecifics but avoid pairing generally, behavioral avoidance effectively reduces disease risk, but at levels that concomitantly compromise host social structure.

### Evolution of Behavioral Resistance

We next examined the evolutionary dynamics in a population with genetic variants that do (*X*_2_) and do not (*X*_l_) avoid disease. When behavioral resistance was through specific avoidance of infected individuals and costs were fixed, *X*_l_ and *X*_2_ could stably coexist when levels of avoidance by *X*_2_ were high, and over an increasing range of costs, including very high costs to the avoider (>50% reduction in birth rate) at high levels of avoidance (Fig. 2a). When behavioral resistance was through general avoidance, the same overall pattern emerged, but the spread of resistance required much higher levels of avoidance, and the coexistence of *X*_l_ and *X*_2_ was only possible under extreme levels of avoidance, although still over a wide range of costs (Fig. 2b).

**Figure 2:**
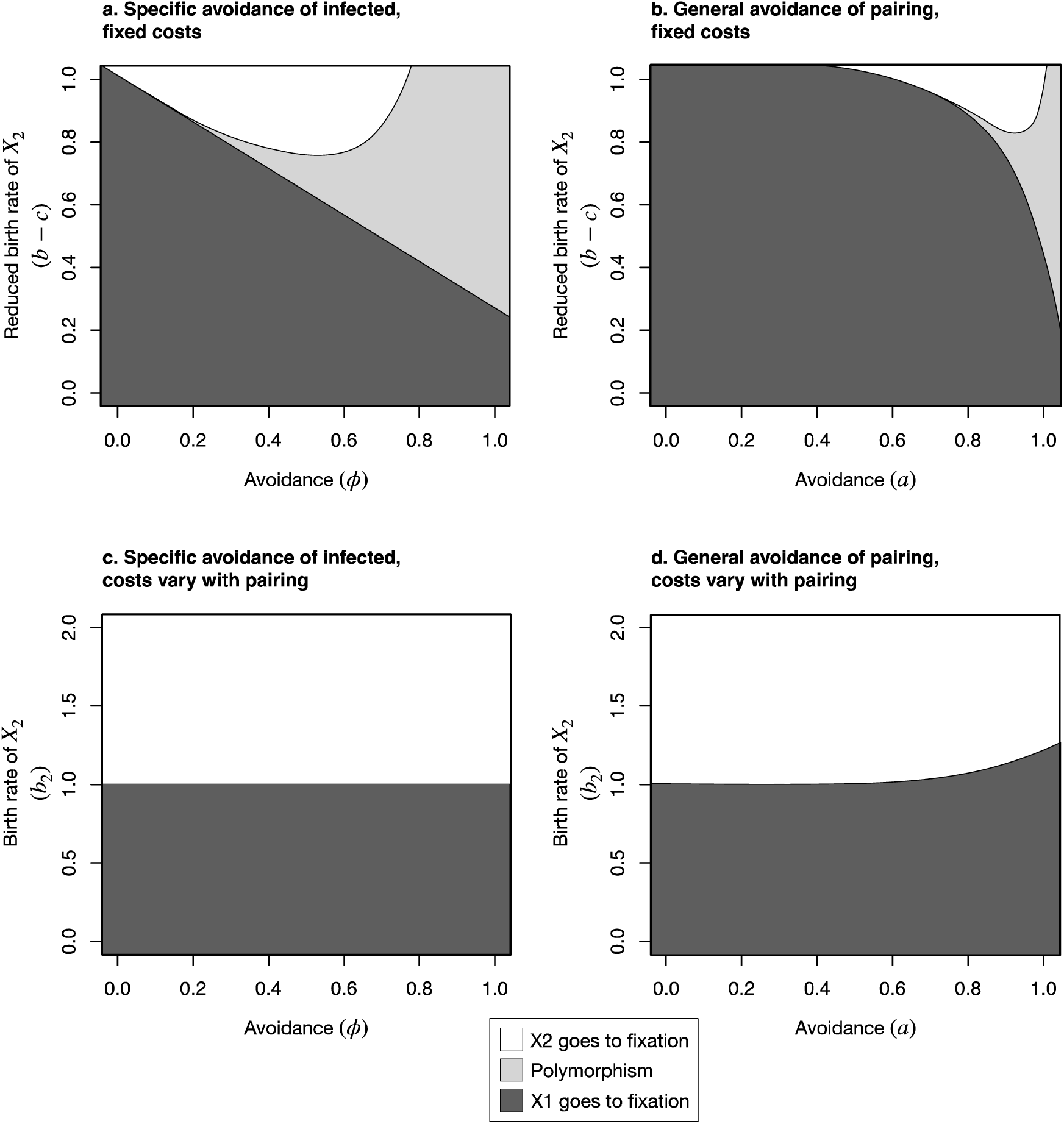
Shaded areas represent equilibrium gene frequency states for the models when the cost and the avoidance strategy of *X*_2_ are varied. *b* = 1, *μ* = 0.2, *δ* = 1, *ρ* = 1, *v* = 0.3, *k* = 0.01.

When costs were incurred as a consequence of not being in a group, in the case of specific avoidance, the benefits of reduced disease risk balanced the costs of lost social interactions, such that *X*_2_ went to fixation when its birth rate was higher than *X*_l_ (Fig. 2c). When avoidance was general, *X*_l_ could even sometimes reach fixation when *X*_2_ had a higher birth rate, because at high rates of general avoidance, reductions in disease risk were not substantial enough to compensate for the loss of social contacts (Fig. 2d). In both cases, when costs were linearly dependent on the frequency of individuals in pairs, polymorphism between *X*_l_ and *X*_2_ was not possible.

## Discussion

Our results show that the dynamics of behavioral resistance can differ from physiological or biochemical resistance evolution depending on the nature of the social behavior and whether the costs are fixed or depend on sociality. As expected, avoidance of social interactions with diseased individuals resulted in reductions of disease prevalence, and this was more effective when there was specific avoidance of diseased individuals, as opposed to general avoidance of social interactions. High levels of specific avoidance result in the full preservation of social structure when hosts extirpate the disease through behavioral mechanisms, whereas at levels of general avoidance that prevent disease spread, social structure is harder to maintain. When there is genetic variation in avoidance levels, the spread of genotypes that avoid group formation depends on the type, level, and nature of the costs of avoidance. When avoidance is specific and costs are fixed, the outcomes are identical to those for physiological resistance evolution, including the counterintuitive outcome that stable genetic polymorphism is more likely when resistance is extreme and costs are large rather than small [5,28]. However, when the costs represent the loss of benefits of group living itself, genetic variation in resistance is much harder to maintain. The possibility of stable genetic variation in behavioral resistance suggests not only that mixed avoidance strategies may represent stable states, but also that genetic differences may be at least partially responsible for individual differences in parasite avoidance in many species, including humans [34,35].

The differences we observe between specific and general avoidance highlight the importance of public health interventions like testing, isolating positive cases, and contact tracing for controlling an outbreak of an emergent disease for which asymptomatic transmission is possible, as in COVID-19 [36]. The maintenance of a polymorphic state under some conditions suggests that if avoidance behavior could be performed flexibly in our simple example, a mixed strategy of social distancing that allows for some social interaction might strike a balance between the costs and benefits. If infection is undetectable, a heavier emphasis on isolation and avoidance would be required for such a mixed strategy to work.

To dissect the basic differences between behavioral and physiological resistance we have deliberately kept the models simple. Future application to any specific host-pathogen context would require more complexity in the temporal and social structure of the interactions. For example, in larger groups, transmission within groups and movement between groups would be possible, and behavioral resistance strategies could be more diverse. Additional models could also examine the effect of mortality, rather than reproduction, costs. Consistent with previous research, this simple model highlights trade-offs between the benefits of reducing disease risk and the costs of foregoing other opportunities, whether nutritional [26], reproductive [27], or in the case of our model, social.

Behavioral and physiological resistance are not separate phenomena but likely interact, with behavioral effects being antecedent to physiological resistance, similar to a two-step infection process [37]. In such situations, genetic associations can arise between genes determining resistance, even without any direct physiological interaction. Physiological and behavioral defenses against parasites might also trade-off with one another. For example, house finches that avoid sick conspecifics invest less in immune defenses [22].

A genetic basis for parasite avoidance behaviors has support from knockout experiments in laboratory mice [38] and selective breeding in livestock [39]. There is also direct evidence of genetic polymorphism in social behavior in halictid bees [40]. Behavioral resistance can thus be innate, as we model it, or learned through prior exposure [41–43]. How dynamics of learned resistance differ from innate is a rich direction for future research. Together these responses represent a suite of psychological and cognitive mechanisms that psychologists have termed the “behavioral immune system” [44], and our study shows that how this metaphor translates to dynamics of behavioral resistance merits further examination.

## Code and Supporting Materials

All code and derivations can be found in the Supplementary Materials.

## Authors’ Contributions

CRA and JA conceived the project and derived the equations together. CRA carried out the simulations and drafted the manuscript. JA provided critical input on the simulations and revisions to the manuscript. All authors gave final approval for publication and agree to be held accountable for the work performed therein.

