## Supplementary Materials S1 for "Evolution of behavioral resistance in host-pathogen systems"

**Supplementary Information – S1: Equations**

Table S1. Mathematical symbols used in the text

| Symbol | Description |
| --- | --- |
| $X_{1}$ | Number of susceptible individuals, non-avoiding genotype |
| $X_{2}$ | Number of susceptible individuals, avoiding genotype |
| $Y$ | Number of infected individuals (in some cases, $Y_{1}$ and $Y_{2}$ used to distinguish $X_{1}$ and $X_{2}$ that have become infected) |
| $N$ | Total population size in a given time-step. Constant within a time-step, changes due to births and deaths from one time-step to the next |
| $T$ | Group size (= 2 for the purposes of this study) |
| $G$ | Number of groups (in case of general avoidance $G_{1}$= groups that include $X_{1},$ $G_{2}$ = groups that include $X_{2})$ |
| $S$ | Number of singletons |
| $\rho$ | Rate at which groups are formed (per singleton per time-step) |
| $\upsilon$ | Rate at which groups dissociate (per group per time-step) |
| $a$ | General avoidance parameter: amount by which rate of group formation is decreased in avoider genotypes ($a=0$ in case of specific avoidance) |
| $\phi$ | Specific avoidance parameter: amount by which group formation is decreased if group contains a diseased individual. ($\phi=0$in case of general avoidance) |
| $b$ | Birth rate: number of new individuals born and surviving to adulthood (per existing individual per time-step; when costs are social, $b_{1}$ and $b_{2}$ refer to birth rates of $X_{1}$ and $X_{2}$ respectively) |
| $c$ | Fixed cost of resistance: reduction in birth rate of avoiding individuals |
| $\delta$ | Transmission rate per time-step given a contact |
| $\mu$ | Mortality rate per time-step |
| $R_{0}$ | Basic reproduction number of the pathogen: number of new infections per existing infection, when disease is very rare |

We model the social behavior of the host in a population of individuals that enter into groups or remain as singletons. We assign S as the number of singletons and G the number of groups of size T. The total population size in a given time-step of the model is the sum of singletons and individuals in groups, $N=S+TG$. We assume group formation occurs rapidly, reaching equilibrium within each time-step, prior to disease transmission, and prior to the demographic processes of birth and death. We assume group formation occurs rapidly, and reaches equilibrium within each time-step. Once groups are formed, disease transmission is only possible within groups. Throughout, we assume a large population and deterministic dynamics.

We examine the simplest case of group association, the formation of pairs, i.e. $T=2$. We assume two processes for pair formation. First, pairs could form at a constant frequency.

**If encounter rate is at a constant frequency (corresponding to frequency-dependent transmission)**

The differential equations for number of groups and singletons are given as:

| $\frac{dG}{dt}=\rho S-\upsilon G$ | (SE1) |
| --- | --- |

| $\frac{dS}{dt}=T(\upsilon G-\rho S)$ | (SE2) |
| --- | --- |

By setting $\frac{dS}{dt}$ and $\frac{dG}{dt}$to zero, we can calculate that at equilibrium, the ratio of groups to singletons, $\frac{G}{S}=\frac{\rho}{\upsilon} .$The frequency of individuals in groups, or $\frac{TG}{N}$, can be calculated from this equation as $\frac{\frac{\rho}{\upsilon}}{1+\frac{\rho}{\upsilon}}$ . Thus, the number of groups at equilibrium under frequency-dependent pair encounter is:

| $G=\left( \frac{\frac{\rho}{\upsilon}}{1+\frac{\rho}{\upsilon}} \right)\left( \frac{N}{T} \right)$ | (SE3) |
| --- | --- |

Number of pairs of X1 (if all the individuals in the population were X1):

| $G_{1}=\left( \frac{\frac{\rho}{\upsilon}}{1+\frac{\rho}{\upsilon}} \right)\left( \frac{N}{T} \right)$ | (SE4) |
| --- | --- |

Number of pairs of X2 (if all individuals in the population were X2):

| $G_{2}=\left( \frac{\frac{\rho-a}{\upsilon}}{1+\frac{\rho-a}{\upsilon}} \right)\left( \frac{N}{T} \right)$ | (SE5) |
| --- | --- |

Second, we consider that pairs could encounter one another at higher rates as the density of singletons increases.

**If encounter rate is dependent on the density of singletons (corresponding to density-dependent transmission)**

Such a scenario could be represented by a number of functions, but for the sake of simplicity, we calculate the rate at which groups encounter one another as a function of the number of singletons, $\rho S$. Thus, as more singletons exist in the population, group formation increases.

This gives us the following differential equations:

| $\frac{dG}{dt}=\rho S^{2}-\upsilon G$ | (SE6) |
| --- | --- |

| $\frac{dS}{dt}=T(\upsilon G-\rho S^{2})$ | (SE7) |
| --- | --- |

At equilibrium, $\frac{G}{S^{2}}=\frac{\rho}{\upsilon}$. We can substitute $G=\frac{N-S}{T}$ into this equation, which can be reduced to $T\rho S^{2}+\upsilon S-\upsilon N= 0$. Using the quadratic formula, we can calculate that

| $S=\frac{-\upsilon+ \sqrt{\mu^{2}+4T\rho\upsilon N}}{2T\rho}$ | (SE8) |
| --- | --- |

And the number of pairs at equilibrium can be calculated as follows.

| $G=\frac{N-S}{T}$ | (SE9) |
| --- | --- |

Thus, the number of pairs of $X_{1}$ (if all the individuals in the population were $X_{1}$) is

| $G_{1}=N-\frac{-\upsilon+\sqrt{\upsilon^{2}+4T\rho\upsilon N}}{2T^{2}\rho}$ | (SE10) |
| --- | --- |

And the number of pairs of $X_{2}$ (if all individuals in the population were $X_{2}$) is

| $G_{2}=N-\frac{-\upsilon+\sqrt{\upsilon^{2}+4T\left( \rho-a \right)\upsilon N}}{2T^{2}(\rho-a)}$ | (SE11) |
| --- | --- |

**Number of pairs of each type**

Once we have calculated the equilibrium number of pairs, we can account for transmission of the disease, which only occurs when an $X_{1}$ or $X_{2}$ pairs with a $Y$ individual.

| $P_{X1Y}=2G_{1}\left( \frac{X_{1}}{N} \right)\left( \frac{Y}{N} \right)= \left( \frac{2G_{1}X_{1}Y}{N^{2}} \right)$ | (SE12) |
| --- | --- |

| $P_{X2Y}=\left( 1-\phi\right) 2G_{2}\left( \frac{X_{2}}{N} \right)\left( \frac{Y}{N} \right)= \left( 1-\phi\right)\left( \frac{2G_{2}X_{2}Y}{N^{2}} \right)$ | (SE13) |
| --- | --- |

We can also calculate the total number of $X_{1}$ and $X_{2}$ in pairs, which we use later to calculate the costs that depend on social structure (see below).

Total number of X1 and X2 in pairs:

| $P_{X1} =\frac{{2G}_{1}X_{1}}{N}$ | (SE14) |
| --- | --- |

| $P_{X2} = \frac{{2G}_{2}X_{2}}{N}-\left( \phi\right) 2G_{2}\left( \frac{X_{2}}{N} \right)\left( \frac{Y}{N} \right)$ | (SE15) |
| --- | --- |

where subtracting $\left( \phi\right) 2G_{2}\left( \frac{X_{2}}{N} \right)\left( \frac{Y}{N} \right)$ represents removing the pairs that were NOT formed as a result of specific avoidance of infected individuals.

**Fixed costs**

If costs are fixed, then the following differential equations define the rate of change in the numbers of each type of individual across time-steps:

| $\frac{dX_{1}}{dt} = X_{1}\left( b-kN-\mu\right) -\delta\left( \frac{2G_{1}X_{1}Y}{N_{t}^{2}} \right)$ | (SE16) |
| --- | --- |

| $\frac{dX_{2}}{dt} = X_{2}\left( (b-c)-kN-\mu\right) -\delta\left( 1-\phi\right)\left( \frac{2G_{2}X_{2}Y}{N^{2}} \right)$ | (SE17) |
| --- | --- |

| $\frac{dY}{dt} = \delta\left( \frac{2G_{1}X_{1}Y}{N^{2}} \right) + \delta\left( 1-\phi\right)\left( \frac{2G_{2}X_{2}Y}{N^{2}} \right) -\mu Y$ | (SE18) |
| --- | --- |

**Costs depend on social structure**

If being in a pair provides a benefit, then the following equations define the rate of change of each type of individual:

| $\frac{dX_{1}}{dt} = X_{1}\left( b_{1}\left( 1+\frac{P_{X1}}{X_{1}} \right)-kN-\mu\right) -\delta\left( \frac{2G_{1}X_{1}Y}{N^{2}} \right)$ | (SE19) |
| --- | --- |

| $\frac{dX_{2}}{dt} = X_{2}\left( (b_{2}\left( 1+\frac{P_{X2}}{X_{2}} \right)-kN-\mu\right) -\delta\left( 1-\phi\right)\left( \frac{2G_{2}X_{2}Y}{N^{2}} \right)$ | (SE20) |
| --- | --- |

| $\frac{dY}{dt} = \delta\left( \frac{2G_{1}X_{1}Y}{N^{2}} \right) + \delta\left( 1-\phi\right)\left( \frac{2G_{2}X_{2}Y}{N^{2}} \right) -\mu Y$ | (SE21) |
| --- | --- |

**Transmission term in behavioral vs. physiological resistance**

When all individuals are in pairs, the model of behavioral processes of resistance is identical to a mass action model of mixing behavior. To illustrate this, using the $X_{1}$ genotype as an example, the transmission term is classically represented by $\beta X_{1}\left( \frac{Y}{N} \right)$. In the present behavioral model, when all individuals are in pairs,$G =\frac{N}{2}$. Substituting this fraction for G in the transmission term for $X_{1}$above gives

| $2\delta\left( \frac{N}{2} \right)\left( \frac{X_{1}}{N} \right)\left( \frac{Y}{N} \right)$ | (SE22) |
| --- | --- |

which simplifies to the equivalent $\delta X_{1}\left( \frac{Y}{N} \right)$. The same follows for transmission to $X_{2}.$

**Calculating** $\boldsymbol{R}_{\boldsymbol{0}}$

$R_{0}$, or the basic reproductive number, is the condition that must be met for new infections to be produced in any time-step of the model. In other words, the differential equation for Y must be greater than 0.

| $0<\frac{dY}{dt}=\frac{2\delta GY}{N^{2}}(X_{1}+X_{2})-\mu Y$ | (SE23) |
| --- | --- |

When Y is rare,$X_{1}+X_{2}\approx N$, which gives

| $0<\frac{2\delta G}{N}-\mu$ | (SE24) |
| --- | --- |

This can be further reduced to the formula for $R_{0}$:

| $1<\frac{2\delta G}{N\mu}=R_{0}$ | (SE25) |
| --- | --- |

This formulation represents the frequency of pairs $\frac{2G}{N}$ multiplied by per contact transmission probability ($\delta$), divided by the background mortality rate ($\mu$).

**Two classes of infected individuals**

We also examined the degree to which our assumption that infected individuals ($Y$) could be grouped together in terms of their behavior affected our results. We thus formulated the equations to make $X_{1}$ individuals become $Y_{1}$ upon infection, and $X_{2}$ become $Y_{2}.$ $Y_{2}$ individuals would thus continue to avoid conspecifics (either generally or diseased specifically), despite their own infection.

If this is the case, then the number of pairs of each type relevant for transmission would be calculated as follows:

| $P_{X1Y1}= \frac{2G_{1}X_{1}Y_{1}}{N^{2}}$ | (SE26) |
| --- | --- |

| $P_{X1Y2}= \frac{2G_{2}X_{1}Y_{2}}{N^{2}}$ | (SE27) |
| --- | --- |

| $P_{X2Y1}=\left( 1-\phi\right)\left( \frac{2G_{2}X_{2}Y_{1}}{N^{2}} \right)$ | (SE28) |
| --- | --- |

| $P_{X2Y2}=\left( 1-\phi\right)\left( \frac{2G_{2}X_{2}Y_{2}}{N^{2}} \right)$ | (SE29) |
| --- | --- |

For the sake of transmission, the only term that is different from the original model (with one class of $Y$) is the pairing between $X_{1}$ and $Y_{2}$ in the case of general avoidance (i.e. this term is calculated from $G_{2}$, which is a function of ($\rho-a$), rather than $G_{1}$ as in the original model).

It follows that the corresponding differential equations are:

| $\frac{dX_{1}}{dt}=X_{1}\left( b-kN-\mu\right)-\delta(P_{X1Y1}+P_{X1Y2})$ | (SE30) |
| --- | --- |

| $\frac{dX_{2}}{dt}=X_{2}\left( (b-c)-kN-\mu\right)-\delta(P_{X2Y1}+P_{X2Y2})$ | (SE31) |
| --- | --- |

| $\frac{dY_{1}}{dt}=\delta\left( P_{X1Y1}+P_{X2Y1} \right)-\mu Y_{1}$ | (SE32) |
| --- | --- |

| $\frac{dY_{2}}{dt}=\delta\left( P_{X1Y2}+P_{X2Y2} \right)-\mu Y_{2}$ | (SE33) |
| --- | --- |

When costs depend on social structure, the equations follow the same format as *SE19-SE21*, except that the frequency of $X_{1}$ in pairs depends in part on the number of pairings with $Y_{2},$ which is a function of $G_{2}$ (rather than of $G_{1}$) in the case of general avoidance.
