## Supplementary Materials S2 for "Evolution of behavioral resistance in host-pathogen systems"

**Supplementary Information – S2: Additional Results**

We present here the results from the case of density-dependent pair encounter, which corresponds to density-dependent transmission.

*Dynamics with No Evolution when Pairing is Density-Dependent*

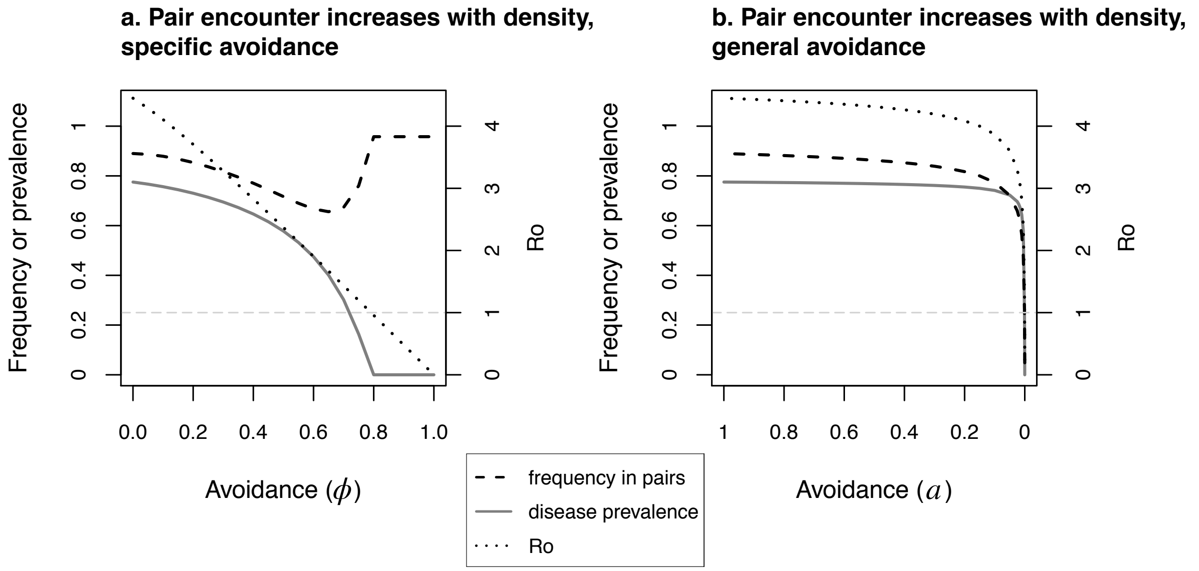


**Figure S1.** Pairing and disease dynamics at equilibrium when only the avoiding genotype $X_2$ is present in the population under different avoidance strategies. The light gray horizontal dotted line represents when the basic reproductive number $R_0 = 1$, below which the disease cannot persist in the population, and above which sustained transmission is possible. $b = 1, \mu = 0.2, \delta = 1, \rho = 1, v = 0.3, k = 0.01$.

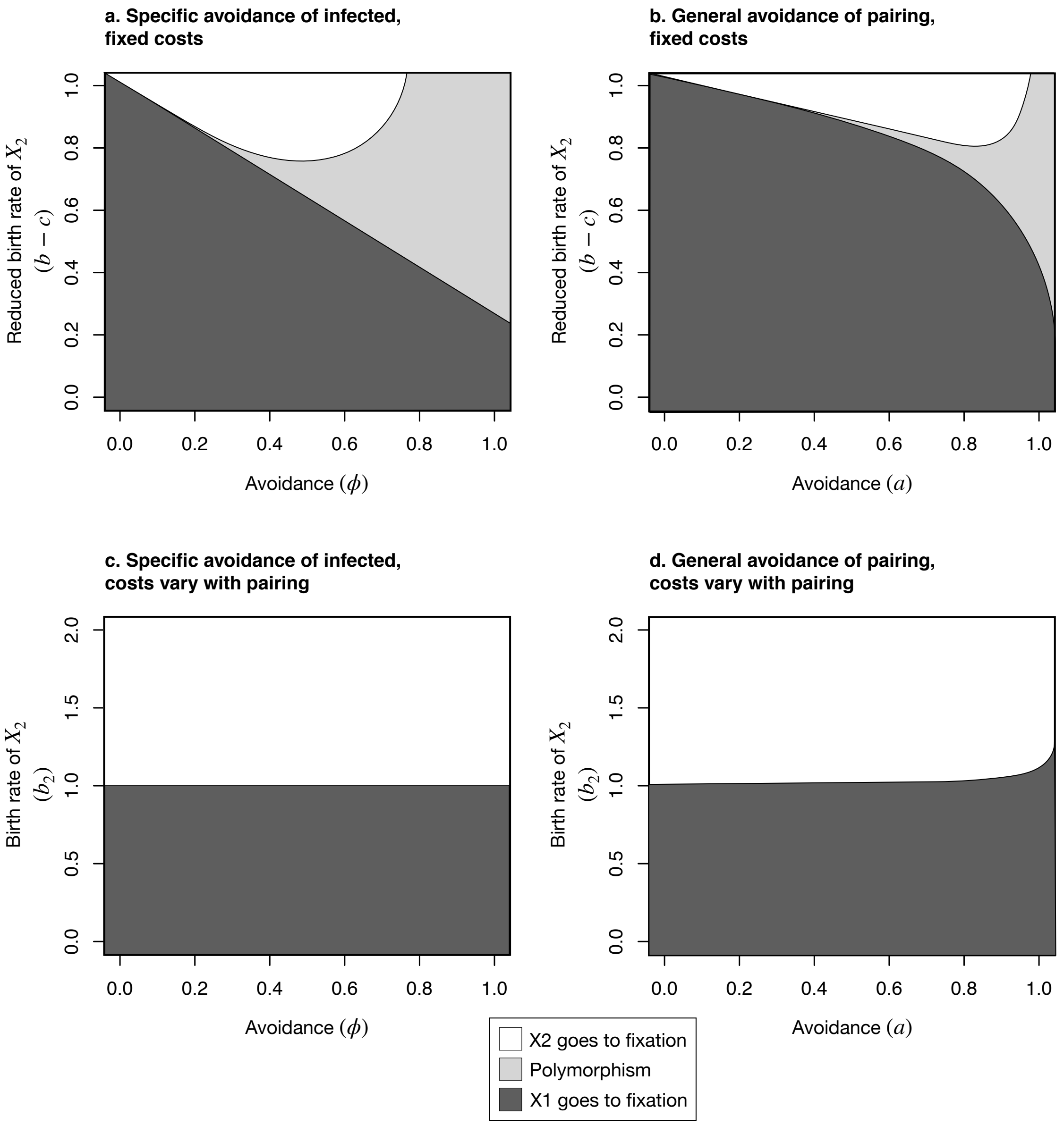


**Figure S2:** Shaded areas represent equilibrium gene frequency states for the models when the cost of $X_2$ and the avoidance strategy of $X_2$ are varied. All plots show results for density dependent encounter rate. $b = 1, \mu = 0.2, \delta = 1, \rho = 1, v = 0.3, k = 0.01$.

*Infected Individuals ($Y_2$) Also Avoid Disease*

We provide results from the case of density-dependent group encounter when infected individuals retain their avoidance phenotype. In other words, when $X_1$ is infected, it becomes $Y_1$, and when $X_2$ is infected, it becomes $Y_2$. We thus assume that $Y_1$ and $Y_2$ have the same avoidance behaviors as their susceptible counterparts. For specific avoidance, the results are identical. For general avoidance the results are qualitatively the same, but the boundaries of the equilibrium states are slightly different.

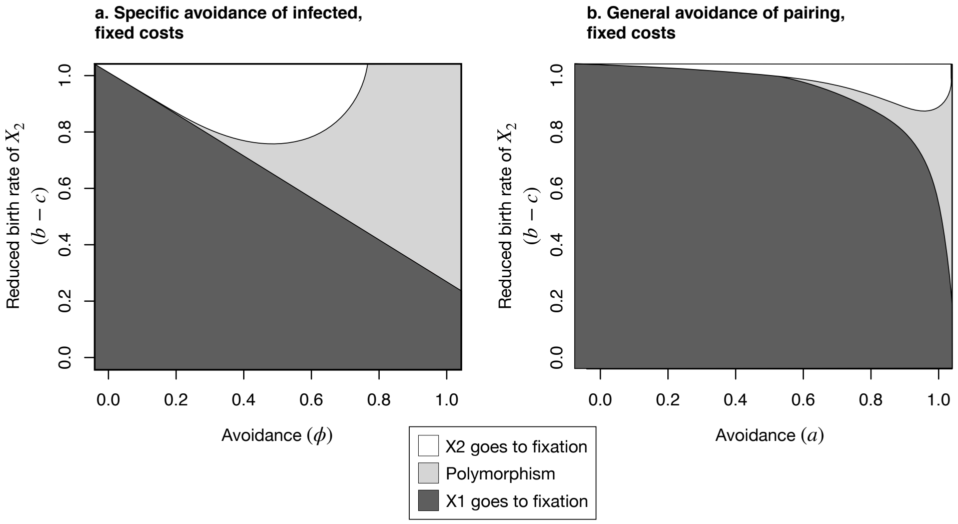


**Figure S3:** Gene frequency results when avoiding and non-avoiding classes of infected individuals ($Y_1$ and $Y_2$) are differentiated in the model. Shaded areas represent equilibrium gene frequency states for the models when the cost of $X_2$ and the avoidance strategy of $X_2$ are varied. Both plots show results for density dependent encounter of pairs. $b = 1, \mu = 0.2, \delta = 1, \rho = 1, v = 0.3, k = 0.01.$
